# The Molecular Evolution of the SARS-COV-2 Spike Protein: Study of Amino Acid Substitutions and Types

**DOI:** 10.1101/2024.07.22.604652

**Authors:** Mike M Moradian, Ania Baghoomian, Michael Sweredoski, Tobias Ivan, Ruan Ramjit

## Abstract

The molecular evolution of SARS-COV-2 has been challenging to predict. Emergence of the Omicron Variant of Concern (VOC) and its sublineages indicated that SARS-COV-2 could evolve more rapidly than previously thought. We analyzed the mutation and amino acid substitution patterns in the spike (S) protein of SARS-COV-2 VOCs to assess how they evolved in response to the selective pressure exerted by both natural immunity and vaccination.

Our results indicate less evolutionary constraint on the first part of the S protein, allowing more amino acid substitutions, especially in the NTD, RBD, and subdomain 1. Omicron lineages introduced mutations in the FP and HR1 domains for the first time. The NTD, subdomain 1, and FP domains allowed more radical amino acid substitutions, followed by RBD and HR1, possibly due to their function. There were up to nine conservative and one radical substitutions in the amino acids interacting with the Human ACE2 receptor in the RBM of the Omicron sublineages only, a remarkable departure from the previous VOCs.

We show that the molecular evolution of SARS-COV-2 S protein was relatively limited up to the Omicron lineage. The selective pressure from previous VOCs and global vaccination potentially accelerated the emergence of the highly transmissible Omicron lineage. This antigenic drift in Omicron is fueled by a high rate of radical amino acid substitutions in the S1 domains, resulting in positive selection with a high potential to change due to adaptive evolution. However, the conservative nature of changes in the RBM may signal a relative stabilization.

## Introduction

The molecular evolution of the coronavirus (SARS-COV-2), which has been responsible for a global pandemic with severe and acute respiratory diseases and syndromes, has been quite remarkable. The SARS-COV-2 has evolved rapidly, resulting in at least 4 significant periods of rapid transmission and spread throughout the world. Although mutations have occurred throughout the entire SARS-COV-2 genome, the mutations in the S gene/spike protein have been instrumental in the transmissibility of the virus. Consequently, several studies have focused on the molecular evolution of the spike protein of the SARS-COV-2, indicating that the spike protein is a variable region (Chayan Roy, 2020, Yuan Huang 2020). They have also reported many non-synonymous mutations in the S gene, which have resulted in both conservative and radical amino acid substitutions, especially in domains such as NTD and RBD. Of course, the acquisition of basic amino acids such as arginine (R) and Histidine (H) in the fusion peptide has also contributed to more efficient fusion of the S protein to the host ACE2 receptors (Kyle Wolf 2022).

The mutations in the S gene and their impact on the spike protein have been different in the major SARS-COV-2 strains. The D614G substitution, which occurred early in the pandemic and gave rise to a more transmittable virus, was one of the first major evolutionary changes that were quickly selected and fixed in the population (Lizhou Zhang 2020). All the subsequent major strains of the SARS-COV-2, including the alpha, delta, and omicron have kept the D614G substitution, making it one of the original/founder substitutions. In contrast to D614G, several mutations have disappeared from the major SARS-COV-2 strains, when one strain took over the previous one as the major strain in the world. For example, mutations Del69/70, Del144, N501Y, and others in alpha were not present in delta and the mutations T19R, Del157/158, P681R, and D950N in delta were not present in omicron. These differences indicated that different lineages of the SARS-COV-2 may have evolved independently over time Volz, E. et al. 2021. While each of the major strains was more transmissible than their predecessor, they had different mutations in their S gene. This could make it challenging to follow, or predict an evolutionary pattern. The S gene’s mutation frequencies have changed over time between each strain, studying these mutation frequencies and their types over time could identify the mutations that have been more effective in the virus’s transmissibility. It could also identify the combination of the mutations in a certain time period that could be directly proportional/responsible for the SARS-COV-2 surges.

Another interesting area of the SARS-COV-2 molecular evolution has been the changes in the amino acids that directly bind/interact with the amino acids on the human ACE2 receptor protein. The substitution N501Y, which was a signature substitution of the alpha strain and has been attributed to this strain’s high transmissibility, is a good example of a substitution in a binding/interacting amino acid on the spike protein (lu Lu 2021). Analysis of the type of amino acid substitutions and their frequency in a certain time period or strain could further identify variable or conserved regions in the spike protein amino acids binding/interacting with the host’s ACE2 receptor. Additionally, the rate of evolution of the SARS-COV-2 was thought to decrease gradually over time before the emergence of the Omicron lineage (shihang Wang 2022). They showed that the coronaviruses have roughly the same mutation rates as the other Flu viruses, Orthomyxoviridae. However, SARS-COV-2 has shown a different and rather unpredictable evolutionary path so far. The emergence of over 30 mutations in the spike protein of Omicron strains Viana, R. et al. Rapid compared to <10 mutations in the previous lineages could be an indication of higher rates of evolution for SARS-COV-2 due to adaptive evolution (Kathryn Kistler & Bedford 2021).

In this study we performed a comprehensive molecular evolution study of the SARS-COV-2 spike protein using the mutation/nucleotide and amino acid substitution patterns. We determined the conserved and variable domains of the spike protein, using radical and conservative amino acid substitution patterns and frequencies.

## RESULTS

### Deviations from expected number of mutations

We predicted the spike protein’s expected proportion of mutations per domain based on the domain length (Figure 1). In the closer VOC linages to Wuhan (i.e., Beta and Gamma), the mutations were concentrated in the NTD, RBD, and subdomain 1 (SD1, the area between RBD and the fusion peptide); all three domains had a higher proportion of mutations than expected. The picture was slightly different in Alpha and Delta, where a higher-than-expected proportion of mutations appeared in the HR1 domain. While in Alpha the proportion of mutations in the RBD was in line or slightly lower than the expectations, a sudden increase in SD1 mutations could have had a compensatory effect on the low mutation frequency in the RBD. However, the emergence of N501Y substitution, which directly interacts with an ACE2 receptor amino acid, along with P681H could have increased the efficiency of the Alpha variant and reduced the need for more RBD mutations. The NTD in Delta has more than twice the expected proportion of mutations, which could have been the reason for Delta’s efficient transmissibility, possibly evading the neutralizing antibodies.

**Figure 1.**
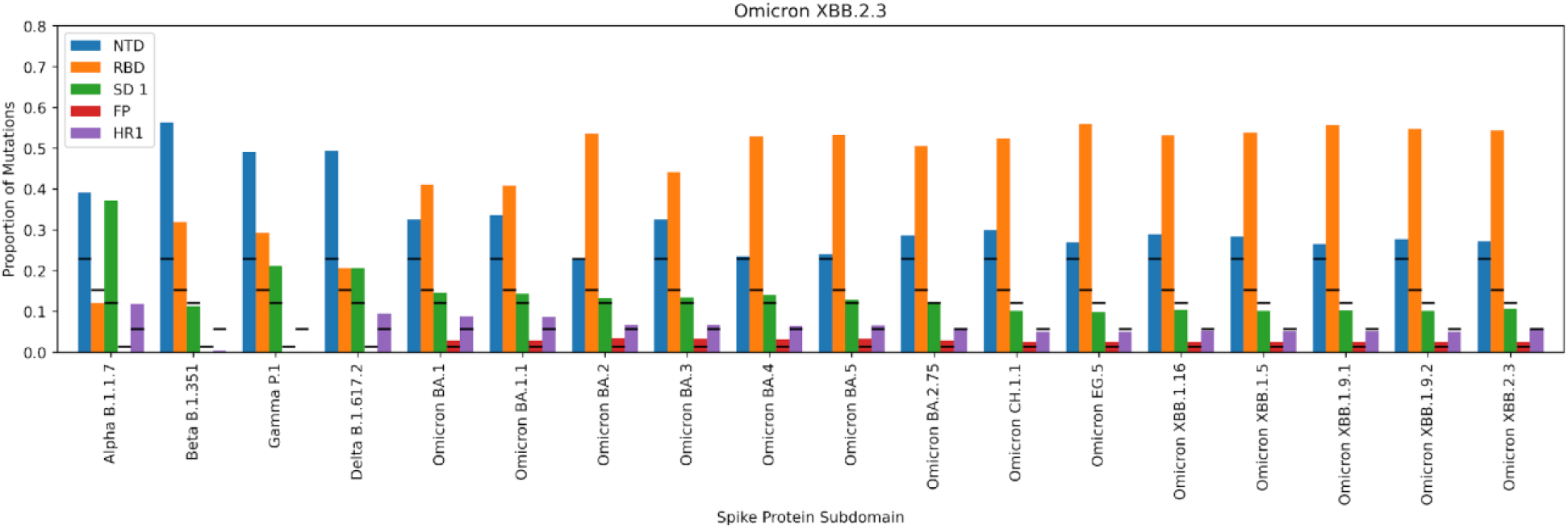
Amino acid substitutions per lineage in Spike protein (black bars are expected proportion of substitutions per domain based on length)

The high proportion of mutations in the Omicron lineage, especially in the RBD is remarkable. For the first time since the pandemic began, in any lineage, there are more than the expected proportion of mutations in the FP domain. Omicron lineages seemingly have a sufficient number of mutations in the NTD to possibly evade the neutralizing antibodies. Furthermore, they have a very high proportion of mutations in the RBD, SD1, FP, and HR1 domains to possibly transmit and infect the host cell more effectively. Also, they have 7-9 mutations in the amino acid positions on the spike protein that interact with the ACE2 receptor amino acids, which could potentially increase the efficiency of their host cell binding and transmission.

### Mutations and Amino Acid Substitutions in the Spike Protein

We analyzed the mutations in the spike protein of the VOCs based on their frequency, types, and patterns in each respective VOC. There were significant differences between the number of mutations amongst the VOCs per domain (NTD p-value: 5.7 × 10^-2, RBD p-value: 1.8 × 10^-9, SBD 1 p-value: 1.8 × 10^-2, Fusion Peptide p-value: 1.1 × 10^-4, and HR1 p-value: 4.3 × 10^-4. While Omicron sublineages had the highest number of mutations, they were not evenly distributed across the S protein (Figure 2). Mutations in Omicron sublineages were concentrated in the RBD (i.e., up to 22 mutations), at least half of which occurred in the RBM. In the NTD, the presence of several amino acid deletions and substitutions indicates several variable positions in this domain. These include amino acid positions 18-27 (Del 25/27 in all Omicrons), positions 67-70 (Del 69/70 in Alpha and BA.2.86), positions 142-158 (Del 144 in Alpha and all Omicrons except CH.1.1, and Del 157/158 in Delta), and the positions 211-215 (Del 212 BA). These deletion events along with several other substitutions indicate that the NTD could tolerate a fair number of changes and may not be under strong evolutionary conservation pressure.

**Figure 2.**
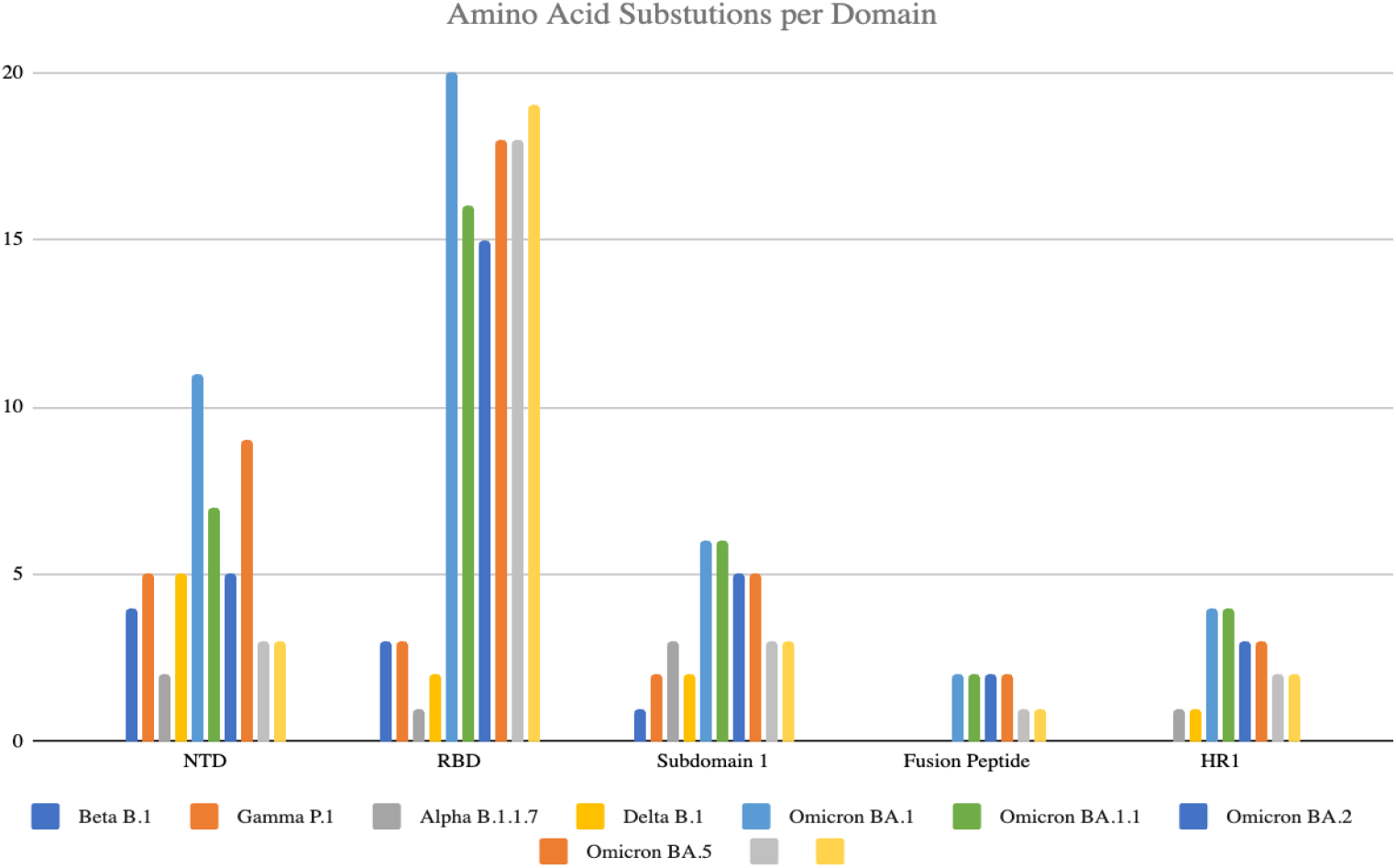
Amino acid substitutions per domain

While the other regions/domains in S protein, such as SD1, FP, and HR1 had fewer amino acid substitutions, the numbers were similarly different between VOCs, with more mutations in the Omicron sublineages.

The main difference between the Omicron lineages and the rest of the VOCs is in the RBD/RBM. Omicron sublineages have up to 22 mutations in this domain while the rest of the VOCs had 9 combined. Only two of the Omicron mutations in the RBM have been previously reported (K417N in Beta and N501Y in Alpha, Beta, and Gamma), the rest are novel. The N501Y substitution, which is considered a key mutation that increases the transmission of the SARS-COV-2 (https://www.nature.com/articles/s41586-021-04245-0) please add this reference hereand is present in almost all the VOCs, was absent in Delta. All Alpha and Omicron lineages (except for BA.2.86) have the P681H substitution in the S1/S2 cleavage site This substitution represents a key difference between the Alpha and Omicron lineages and the Delta and BA.2.86 lineages, which have the substitution P681R.

The amino acid substitution patterns showed interesting differences between the VOCs. The VOCs before Alpha, such as Beta and Gamma, which did not spread as rapidly globally, did not have any amino acid deletion in the NTD, while all the VOCs after alpha had at least 2 and up to 5 amino acid deletions, suggesting that these deletions may have played a potentially significant role in their global transmission.

We also observed more radical amino acid substitutions across all the VOCs in the NTD than conservative substitutions (Figure 3). In contrast, up to the Omicron lineage the RBD had more conservative substitutions than radical substitutions, which were primarily in the RBM region. Omicron sublineages had radical amino acid substitutions throughout the RBD domain with up to half of them being radical in the RBM, suggesting less conservation across this domain. There were more radical than conservative amino acid substitutions in SD1 and the FP domains. In fact, FP had only radical amino acid substitutions and only in the Omicron sublineages. Except for Alpha, the HR1 domain had more conservative than radical amino acid substitutions in all the VOCs. In Alpha, there was only one amino acid substitution (radical), a distinct difference from the other VOCs.

**Figure 3.**
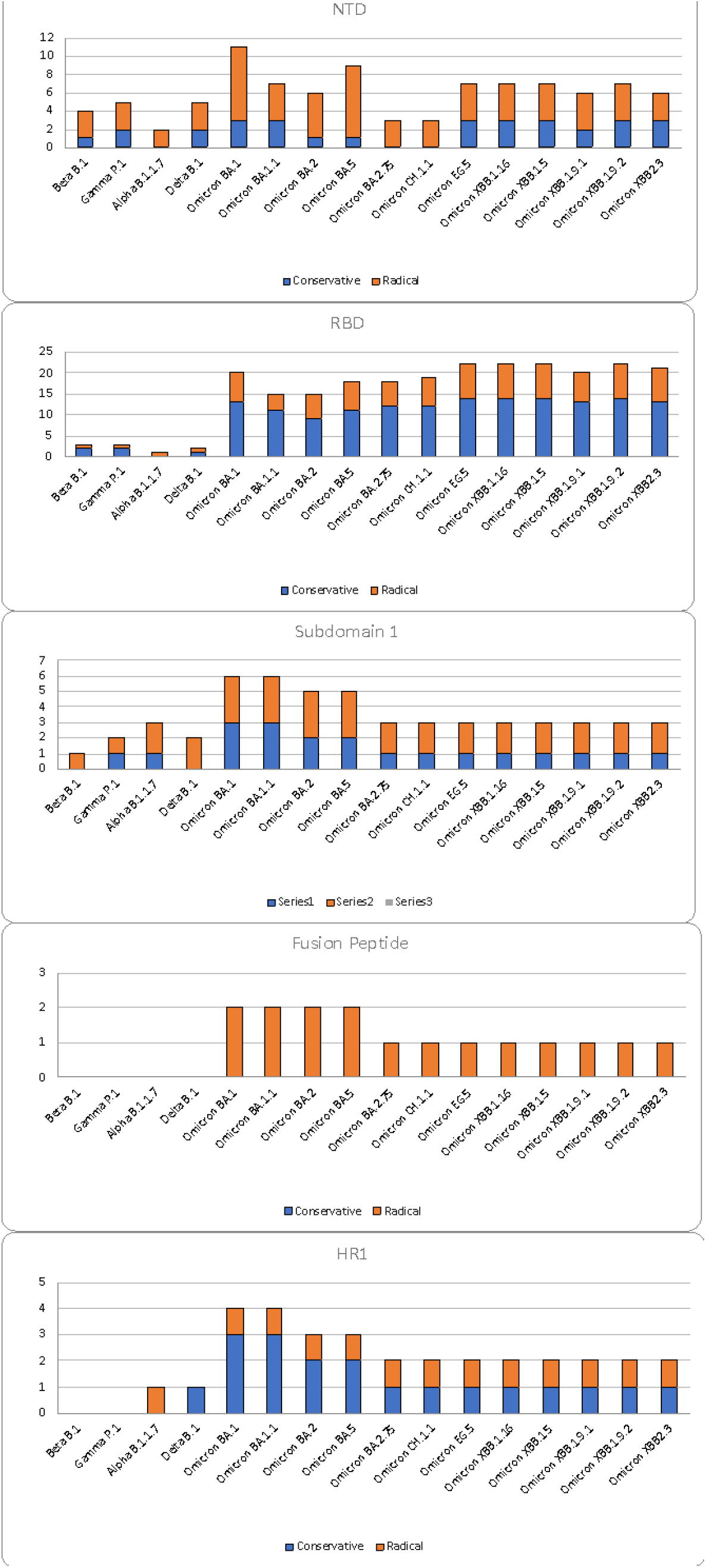
Radical and conservative substitutions per domain in each VOC lineage

Analysis of the substitution patterns in the RBM amino acids, (i.e, R403, K417, N439, G446, G447, Y449, Y453, L455, F456, A475, G476, S477, E484, F486, N487, Y489, P491, Q493, G496, F497, Q498, T500, N501, G502, Y505, Q506) that are believed to be interacting with the human ACE2 receptor (Hocheol Lim and Jun Lan) showed up to nine substitutions in Omicron sublineages, (Omicron BA1.1 had nine substitutions, and the rest of the Omicron sublineages had seven or eight). VOC Delta had no substitution in the interacting amino acids and Alpha had one (N501Y). Beta and Gamma had three substitutions (K417N, E484A, and N501Y), all of which were present in Omicron sublineages. There were seven amino acid substitutions (six conservative and one radical) common in almost all Omicron sublineages, two were present in BA1.1 (G496S), BA.EG.5, and BA.1.9.2 (F456L). All of these substitutions, with the exception of N501Y, were conservative, suggesting limited variability allowed in the RBM.

## Discussion

Analysis of the nucleotide and amino acid substitution patterns and types in the S protein of the SARS-COV-2 allowed us to provide further evidence of the molecular evolution of this virus. The amino acid substitution types indicated that the S protein allows two very different patterns of evolution, a highly variable region starting at the NTD to the HR1 domain and a highly conserved region thereafter. Different studies have already suggested the reasons for such structure (Irwing Jungrais 2021, Yuan Huang, 2020), yet the emergence of the Omicron lineage has changed the molecular evolution of the SARS-COV-2. The domains in the S1 region of the Omicron lineage proved to be highly variable, possibly resulting in an antigenic drift from previous VOCs. The main difference was in the RBD/RBM between the Omicron sublineages and the rest of the VOCs. This region, which contains the amino acid residues that interact with the ACE2 receptor in human cells, has been quite conserved in the previous VOCs, yet introduction of up to 22 amino acid substitutions in Omicron sublineages completely changed that picture. These nonsynonymous mutations, resulting in amino acid substitutions in the RBD indicate an evolutionary response to the pressure from antibodies due to vaccination and prior infections in SARS-COV-2. Two studies presented 26 amino acids in the RBM that they believed interact with the human ACE2 (Hocheol Lim and Jun Lan). All the VOCs before Omicron had between zero and three substitutions, Omicron sublineages have up to nine, meaning about one-third of the presumptive interacting amino acids in the Omicron RBM were mutated/substituted. This adaptation could be the reason for Omicron’s supreme transmissibility.

Since the RBD/RBM amino acids interact with ACE2, radical changes—especially in higher numbers—may be disadvantageous to the SARS-COV-2 transmissibility. The adaptive evolution of the S1 domains in the Omicron lineage was not uniform; all domains, except the RBD, had more radical changes/amino acid substitutions. The relative stability (i.e., more conservative amino acid substitutions) in the RBD could be primarily due to the fact that eight out of nine amino acid substitutions in the RBM, which interact with the human ACE2 protein were conservative; the exception was N501Y. The domination of the conservative amino acid substitutions in the RBM is notable, suggesting that there may be a limit to the antigenic drift in this domain. If RBM does not allow radical changes to its structure, then its adaptive evolution may be stabilizing.

Substitutions at residues L452, F486, and R493, which are targets for neutralizing antibodies in the RBM, have been shown to confer resistance in the Omicron sublineages (Qian Wang et al., 2022). Although most of the amino acid substitutions in the RBM of the Omicron sublineages were conservative, the BA.5 RBM had three radical substitutions (highest among all Omicron sublineages) from a total of nine. This sublineage has also shown substantially more neutralization resistance to sera obtained from vaccinated and boosted individuals (Khan K et al., 2022). If there is any correlation between radical amino acid substitutions in the RBM and antibody evasion, then we may anticipate seeing more of them in the future due to vaccine boosters.

The rates of evolution for SARS-COV-2 have been suggested to be in line with other similar respiratory viruses (Shihang Wang et al). This was true up to the Delta lineage, yet the Omicron lineage proved that perhaps it is early to be able to estimate the evolutionary rate of the SARS-COV-2 accurately. Kistler et al. (reference) presented an analysis where they estimated the rate of S1 mutations to be four times that of the influenza H3N2 virus. With such a high drift rate in the antigenic region of the SARS-CoV-2, the need to create vaccine boosters for the new lineages has become a reality. This could create a vicious cycle by putting additional selective pressure on the virus to adapt and evade the new antibodies, consequently, creating new lineages. However, since the pressure would be on the virus’s transmission machinery the pathogenicity of it could remain unchanged. At least this is what was observed with the emergence of the Omicron lineage.

Analysis of the amino acid substitution types and patterns revealed interesting observations. None of the VOCs before Alpha had deletion(s) in their NTD. Qing et al. (2021), suggested that the deletions in the NTD synergize with D614G substitution to enable ACE2-independent S2’ cleavage, which is believed to assist with transmissibility. Also, while these VOCshad the only radical substitution in an interacting amino acid, i.e., N501Y, which increases the ACE2 binding affinity of the S protein (Starr et al 2020), they missed the basic substitutions in position 681, which is a preferred interacting amino acid by furin (Hoffman M, Klein Weber H, 2020). The absence of these two important changes in early VOCs may have been a factor in their slower/limited spread in the world. Interestingly, N501Y seems to be present with substitution P681H in more transmittable variants of Alpha and Omicron. While N501Y was not present in the Delta sublineage, the presence of substitution P681R could compensate for this missing mutation, creating the basic cleavage motif in the P5 site. Arginine in position 681 confers the greatest increase in cleavage efficiency (Jaimes JA, Millet JK, 2020). This raises an interesting question and a future opportunity to investigate if there is any connection between the amino acid positions 501 and 681 in the S protein binding and fusion to the human cells.

Overall, adaptive evolution seems to be the driving force for the rapid molecular evolution of the S protein in SARS-COV-2 (Kathryn Kistler 2022). Accumulation of the nonsynonymous mutations in the S protein of different lineages has been a good indication of that lineage’s success. Also, specific substitutions may have contributed to the lineages’ success more than the others, these substitutions have been fixed in multiple VOCs through convergent evolution (Martin et al, 2021; Rochman et al 2021). We demonstrated that the remarkable antigenic drift in Omicron sublineages and their effective and persistent presence as the dominant SARS-COV-2 clade for the longest time may be accompanied by the significantly higher rate of radical amino acid substitutions in their S protein. However, these patterns may also point to stabilizing the S protein evolution and less dramatic changes in the spread of the SARS-COV2 throughout the world.

## Materials and Methods

To conduct the analysis of the mutation/nucleotide and amino acid substitution patterns and types in the S protein of the SARS-COV-2 we utilized the platform outbreak.info. “outbreak.info has been enabled by unprecedented global genomic sequencing efforts and we developed every element of the application to fully leverage this capacity; however, genomic sampling varies globally with the vast majority of sequences coming from high income countries; even within well-sampled regions, there is geographic and temporal variation.

“The web application was built using Vue.js v.2.7.14 (https://vuejs.org/), a model–view–view model JavaScript framework that enables the two-way binding of user interface elements and the underlying data allowing the user interface to reflect any changes in underlying data and vice versa. The client-side application uses the high-performance API to perform interactive operations on the database. In addition, All SARS-CoV-2 virus sequence data were provided by the GISAID Global Data Science Initiative and are available at https://gisaid.org/.”

Outbreak.info genomic reports: scalable and dynamic surveillance of SARS-CoV-2 variants and mutations. Karthik Gangavarapu, Alaa Abdel Latif, Julia L. Mullen, Manar Alkuzweny, Emory Hufbauer, Ginger Tsueng, Emily Haag, Mark Zeller, Christine M. Aceves, Karina Zaiets, Marco Cano, Xinghua Zhou, Zhongchao Qian, Rachel Sattler, Nathaniel L. Matteson, Joshua I. Levy, Raphael T. C. Lee, Lucas Freitas, Sebastian Maurer-Stroh, GISAID Core and Curation Team, Marc A. Suchard, Chunlei Wu, Andrew I. Su, Kristian G. Andersen & Laura D. Hughes. *Nature Methods* (2023). doi: 10.1038/s41592-023-01769-3

